# Tumoral CD24 tunes platelets binding and pro-metastatic functions

**DOI:** 10.1101/2025.04.09.648049

**Authors:** Vincent Mittelheisser, Cristina Liboni, Clarisse Mouriaux, Silvia Maria Grazia Trivigno, Louis Bochler, Maria J. Garcia-Leon, Annabel Larnicol, Laetitia Paulen, Tristan Stemmelen, Pierre H. Mangin, Olivier Lefebvre, Jacky G. Goetz

## Abstract

One of the earliest steps of breast cancer metastasis occurs when tumor cells (TCs) disseminate through the bloodstream. There, they interact with several blood components. Among them, platelet favor TC survival and metastatic spread. While the binding of platelet to TC is highly variable, its molecular controls and downstream consequences remain unidentified. Here, we first document that high CD24 expression correlates with increased platelet binding and poorer survival in breast cancer. We further demonstrate that CD24-mediated platelet binding regulates TC cluster formation and resistance to anoikis *in vitro*. Depleting CD24 expression significantly reduces TC metastatic potential by rewiring the metastatic tumor microenvironment (mTME), affecting immune compartments and secreted factors. Overall, our work identifies CD24 as a molecular cue controlling TC-platelet interaction, dictating their metastatic potential. As such, it represents a druggable target to counteract platelet-TC collaboration in metastasis.

## Introduction

Upon cancer progression, tumor cells (TC) disseminate through the bloodstream to colonize distant organs and form life-threatening metastases. During their intravascular journey, circulating TC (CTC) face several hostile forces that are detrimental to their survival (1). Once their entered the bloodstream, CTC rapidly bind, activate and aggregate circulating platelets (2,3). Platelets provide a physical shield to CTC protecting them from destruction by shear forces (4) and by patrolling immune cells (5). In addition, this interaction favors the subsequent TC adhesion and extravasation at distant sites (6,7), facilitating metastatic foci formation. Interestingly, we recently demonstrate that TC interact with platelet in phenotypically distinct ways, which is variable across cellular models and cancer types (8). By inducing a transient thrombocytopenia at the time of TC tail vein injection, we next reveal that high-binding TC rely on interactions with platelet to achieve successful early lung seeding, whereas low-binding TC seeding is independent of the presence of platelet (8). Yet, the molecular determinants of differential platelet binding abilities by TC remain largely unknown.

Building on such limitations, we designed a study aiming at interrogating which molecular cues are responsible for the high *versus* low platelets binding phenotype of TC. Through the examination of publicly available datasets, we found that CD24 expression was significantly increased in TC that highly bind platelets. This is in line with several studies that have reported CD24 as broadly overexpressed in numerous human malignancies, both solid and hematological (9). Interestingly, its expression varies upon cancer progression and is usually tied with a more aggressive course of the disease and poorer survival rate (10,11). Mechanistically, CD24 is a glycosylphosphatidylinositol-anchored membrane protein functioning as an adhesion molecule for P-selectin expressed both on the endothelium and on activated platelets (12,13). Its expression on TC correlates with higher P-selectin-dependent arrest in the lung (14,15) and, via binding to tumor-educated platelets, induces tumor vascularization and growth (16). Yet, the role of TC-expressed CD24 on platelet binding and subsequent metastatic fitness remains unexplored.

In this study, we genetically depleted CD24 expression in breast cancer TC with a shRNA approach and first demonstrated that CD24, in part, tunes TC abilities to bind platelet. We also showed that CD24-mediated platelet binding was crucial for TC pro-metastatic clustering and resistance to anoikis. *In vivo*, shCD24 cells demonstrated a reduced metastatic burden compared to parental cells together with a profound rewiring of the tumor immune microenvironment (TiME). As the relative contribution of endothelial *versus* platelet P-selectin to TC arrest in the lung still remains unclear (14,15), we subjected mice to a short thrombocytopenia prior TC injection. Platelet-depleted mice presented an additional reduction of metastatic burden, corroborating the concept according to which CD24-mediated TC binding to platelet underlies TC seeding and subsequent metastatic outgrowth. Altogether, our study demonstrates that CD24 is a druggable target for preventing metastatic progression as, among others, it favors the binding and function of pro-metastatic platelet.

## Results

### Platelet binding to TC scales with poor prognosis CD24 expression levels

To dissect the molecular clutch controlling platelets binding to TC in the context of BCa, we screened publicly available databases of RNA-sequencing and assess the expression of known receptors involved in TC binding to platelet (12,17–22) (**Fig.1A**). Based on the findings of our previous work, we selected the murine metastatic triple negative breast cancer (TNBC) 4T1 cells as high platelets binding phenotype and compared them to B16F10 (murine melanoma) cells that we previously demonstrated as low platelets binders (8). Besides, to investigate the expression of platelet receptors during metastatic progression, we further evaluated the 67NR cells, which are the non-metastatic isogenic counterpart of 4T1 (23) (**Fig.1A**). By analyzing the differential expression of the identified platelet receptors genes, we observed that P-selectin ligands *CD24* and *Podocalyxin* (PODXL) were solely overexpressed in 4T1 cells compared to 67NR cells and B16F10 cells, suggesting their implication in both high-platelet biding phenotype and metastatic progression (**Fig.1B**). To confirm the relevance of *CD24* or *PODXL* in human breast cancer, we examined their expression in breast cancer patients by exploring RNA-sequencing data from The Cancer Genome Atlas (TCGA) BRCA dataset (**Fig.1C**). Interestingly, *CD24* expression was significantly increased in human breast cancer tissues compared to healthy breast tissues (**Fig.S1A**). Furthermore, in depth analysis of breast cancer PAM50 subtypes revealed that *CD24* expression was mostly increased in TNBC and HER2-amplified patients (**Fig.S1A**). Yet, when stratified, only TNBC patients presenting high *CD24* expression level had a significantly poorer overall survival (**Fig.S1B**). More particularly, *CD24* expression was increased in TNBC patients (Miller cohort) during local lymph node (LN) invasion when compared to their primary tumor (**Fig.S1C**), underscoring its role in breast cancer cells metastatic abilities. When patients were stratified in the TCGA BRCA dataset based on their LN invasion (LN^neg^ vs LN^pos^), we further highlighted that high *CD24* expression led to poorer overall survival in patients presenting lymph node invasion as compared to those without (**Fig.S1D**). In line with these results, high *CD24* level and lymph node infiltration led to a higher risk of disease recurrence (**Fig.S1E**). Conversely, *PODXL* expression was significantly decreased in human breast cancer cells compared to healthy tissue and widespread in several other clusters within the tumor microenvironment (TME) (**Fig.S1F-G**). Moreover, *PODXL* expression had no impact on patients’ overall survival (**Fig.S1H**). Together, these results suggested that *CD24* expression correlated with *i*) high platelet-binding to TC and *ii*) with breast cancer cells metastatic abilities and overall survival. Hence, we decided to further evaluate CD24 role in platelets binding to TCs.

**Figure 1.**
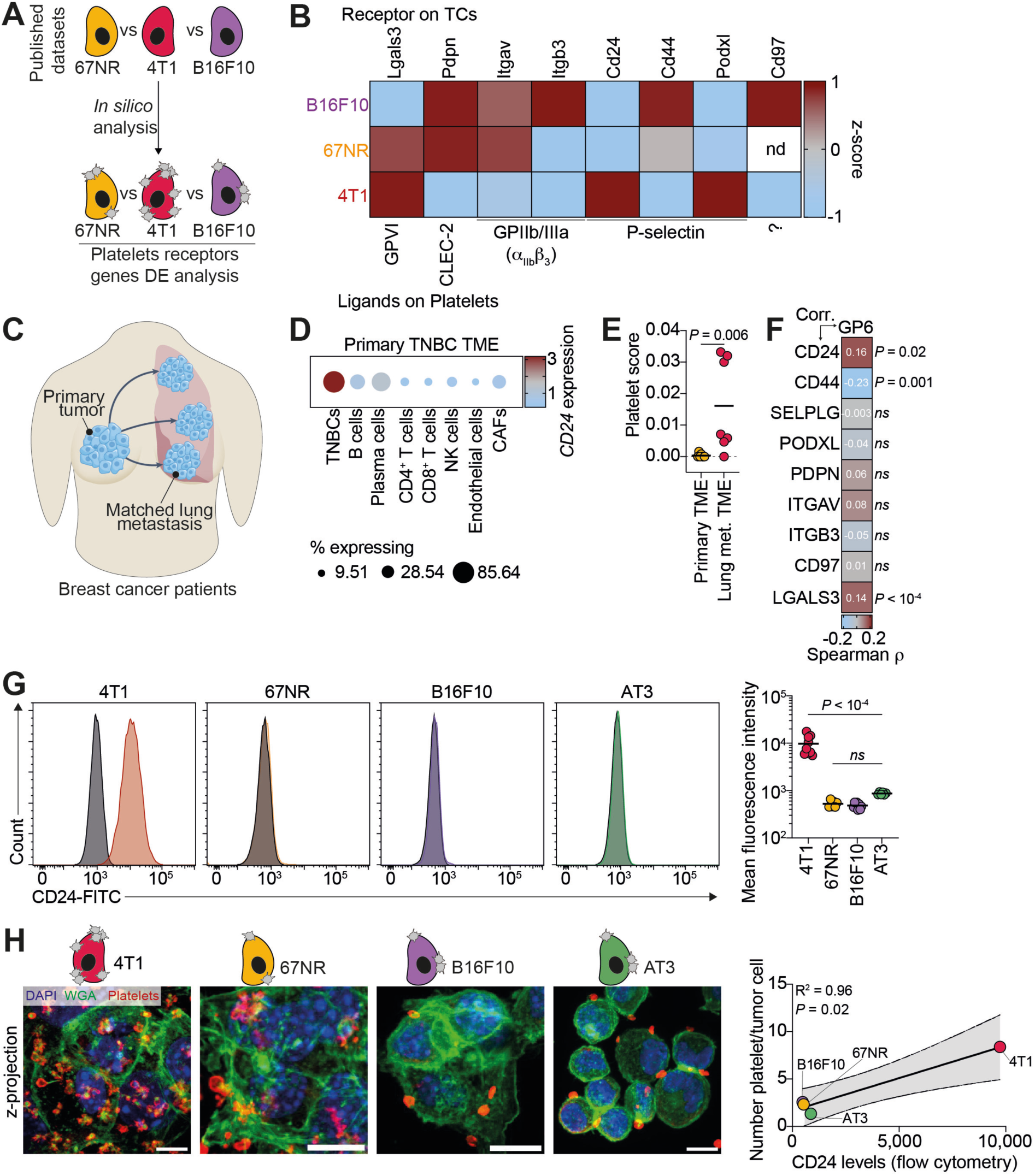
Platelet binding to TC scales with poor prognosis CD24 expression levels. **A**. Infographics illustrating the pipeline of the *in silico* analysis. **B**. Heat-map showing Z-score values for RNA expression of platelet-cognate receptor on tumor cell (TC). Receptors on TC are represented on the top, while cognate receptors on platelet are represented on the bottom. TC analyzed are listed on the left. **C**. Infographics illustrating breast cancer patients’ samples. **D**. Bubble plot displaying *CD24* gene expression at the single-cell level across different cell types in the triple-negative breast cancer (TNBC) primary tumor microenvironment (TME). Bubble size represents percentage of *CD24*-expressing cells and color represent mean *CD24* expression. CAFs: Cancer Associated Fibroblasts. **E**. Platelet score value evaluating platelet infiltration in primary TNBC and their matched lung metastatic TME. Data are representative of 7 different patients from 2 independent clinical studies. Mann-Whitney test was applied after assessment of gaussian data distribution by Shapiro-Wilk test. **F**. Correlation between *GP6* and *CD24*, *CD44*, *SELPLG*, *PODXL*, *PDPN*, *ITGAV*, *ITGB3*, *CD97*, or *LGALS3* expression levels (Log_2_ TPM) in the TNBC subset of the TCGA BRCA cohort (*n* = 191 patients). Color represents the value of Spearman ρ per correlation (legend on the bottom). **G**. Flow cytometry assessment of CD24 expression in selected TC. Left: Representative histograms of the relative expression levels of CD24-FITC. Isotype signal is represented in black in each graph. Right: Quantification of the mean fluorescent intensity signal for CD24-FITC. Data are presentative of between 6 to 9 independent experiments. One-way ANOVA with original FDR method of Benjamini-Hochberg was applied after assessment of gaussian distribution by Shapiro-Wilk test. **H**. *In vitro* platelet-TC interaction. Left: Representative confocal micrographs of platelet-TC binding. In green Wheat Germ Agglutinin (WGA)-iFluor488, in red platelet RAM.1 (Alexa Fluor 647), and in blue nuclei (DAPI). Scale bar = 10µm. Right: Pearson correlation between the number of platelets bound to tumor cell and mean CD24 expression levels on tumor cells – 4 xy pairs; R^2^ = 0.96 – 95% confidence interval [0.31-0.99]. Number of platelets interacting with tumor cell were quantified from 10 fields of view from 2 independent experiments. MeanCD24 expression levels quantified by flow cytometry from G panel were used.

Building on these observations, we next thoroughly investigated *CD24* expression within the TME cell populations analyzing single-cell RNA-sequencing data from 5 primary TNBC samples (24). Tumor cells exhibited robust expression of *CD24*, whereas its expression was weak in all other main TME cell clusters, thus illustrating the potential of CD24 as a tumor-specific target (**Fig.1D**). Since CD24 is associated with lymph node invasion, we further investigated the cellular heterogeneity between the primary TNBC TME and its matched metastatic TME. Focusing on lung metastases, a major site of TNBC spread (25), we performed xCell analysis on primary TNBC and matched lung metastatic lesions (**Fig.1C**) from the Stavanger (3 primary-metastasis pairs) (26) and the University of North Carolina Rapid Autopsy Program (RAP) (4 primary-metastasis pairs) (27) cohorts (**Table S1**). Interestingly, although immune and stromal scores were similar between primary TNBC and their matched lung metastases, we observed a notable increase in platelet infiltration in lung metastatic nodules compared to primary tumors (**Fig.1E**). This platelet infiltration (assessed by *GP6* expression which is restricted to megakaryocytic lineage (28)) within the TME significantly positively correlated with *CD24* expression, but not with other platelet receptors like, *CD44*, *SELPLG*, *PODXL*, *PDPN*, *ITGAV*, *ITGB3* or *CD97* with the exception of galectin-3 (*LGALS3*), the prototypical GPVI receptor (**Fig.1F**).To further validate the translatability of our transcriptomic observations at the protein level, we conducted flow cytometry analyses, which confirmed, in agreement with the *in-silico* analysis, robust CD24 protein expression in 4T1 cells and minimal expression at the surface of 67NR and B16F10 cells (**Fig.1G**). We further extended our investigations by probing CD24 expression in the additional low-platelets binding AT3 cell line (8). As expected, CD24 presented a significantly lower expression at their surface compared to high platelets binding 4T1 cells (**Fig.1G**). Building on our previous study (8), we highlighted a significant positive correlation (R^2^ = 0.99) between CD24 surface expression levels on murine tumor cells (**Fig.S1I**) and the number of platelets bound per cell as seen by scanning electron microscopy. In addition, a second 2D model of platelets binding to tumor cells confirmed the significant correlation (R^2^ = 0.98) between the CD24 surface expression levels and the number of platelets bound per cells as observed by immunofluorescence (**Fig.1H**). Altogether, these results identified CD24 as a new TCs specific rheostat regulating platelet binding and thus pro-metastatic functions.

### CD24 controls the binding of platelet impacting survival and clustering potential of TC

To functionally investigate the role of CD24-mediated platelet binding in cancer progression, we reduced CD24 expression via a shRNA approach in the 4T1 cell line (**Fig.2A**). We first validated the successful knock-down of CD24 at the genomic (RT-qPCR) and protein level (flow cytometry) in the 4T1 shCD24 cells compared to their control counterpart (4T1 shScramble) (**Fig.2B-C**). Of note, CD24 expression levels were similar between 4T1 shScramble and their native counterparts (**Fig.S2A**). These same cell lines were assessed for their ability to bind platelets *in vitro*. In this setting, we observed that 4T1 shCD24 displayed significantly reduced ability to bind platelets (**Fig.2D**), reaching levels similar to “low-binding” cells (**Fig.S2B**). As platelet activation is deeply linked to loss of circular shape (29), we next assessed platelet morphology. In addition, to their increased number, platelets showed a decrease in circularity index when bound to 4T1 shScramble cells compared to 4T1 shCD24 cells (**Fig.2D**) or to CD24^neg^ 67NR, B16F10 and AT3 cells (**Fig.S2C**), suggesting that CD24 expression by TC promoted the binding of activated platelet. We next assessed the ability of platelet to promote the clustering of TC (**Fig.2E**), a step known to promote seeding and metastases formation (30). We previously showed that platelets favored the clustering of 4T1 cells but not of B16F10 cells, in line with their platelet binding profile (8). In agreement with this, CD24 knock-down significantly reduced 4T1 cells clustering number and size (**Fig.2E**). Pro-metastatic clustering impairment was also associated with a reduction of resistance of 4T1 shCD24 cells to anoikis (**Fig.2F**). Overall, these results show that CD24 tunes platelet binding to TC and likely contributing to the pro-metastatic function of platelet, affecting TC clustering and fitness.

**Figure 2.**
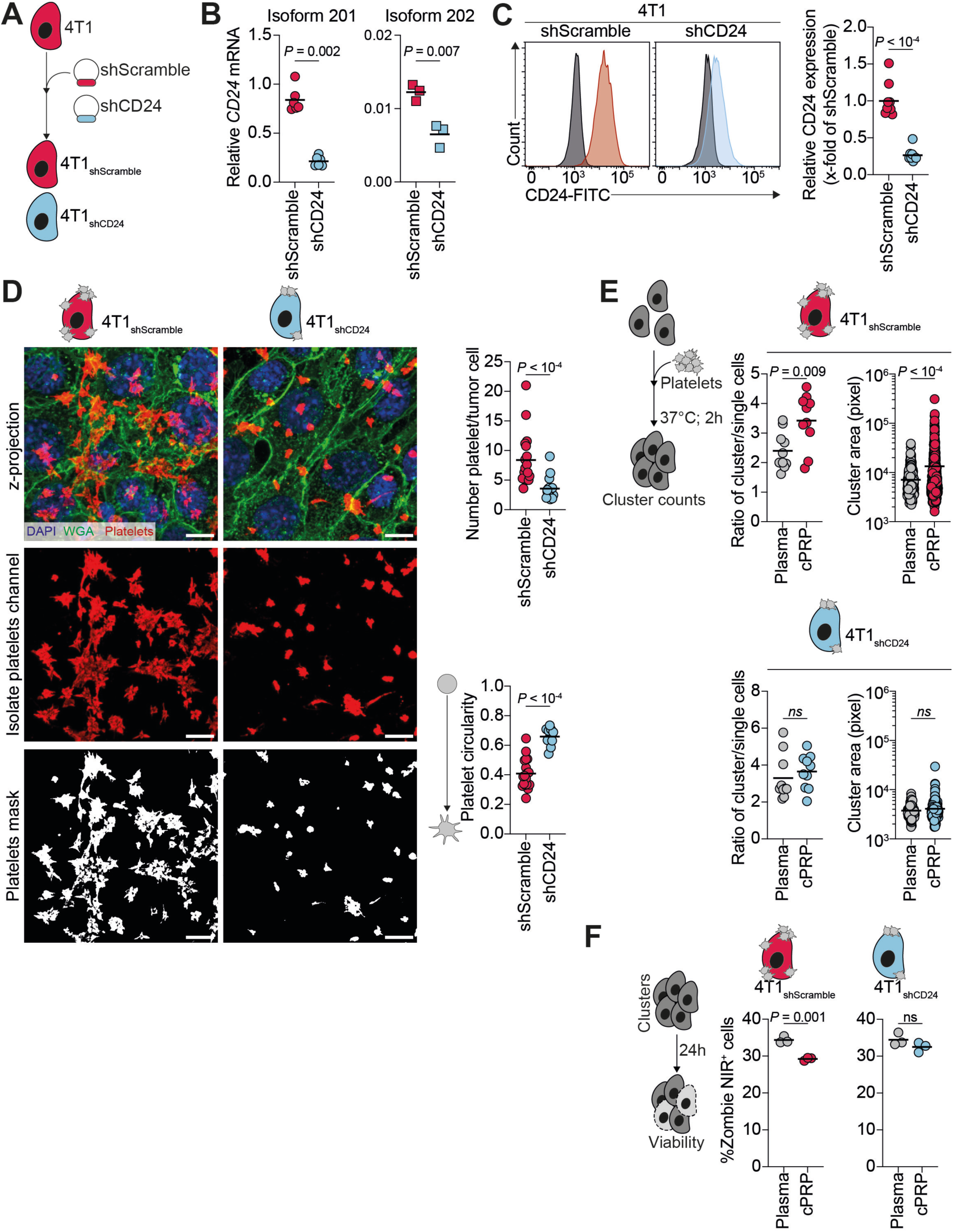
CD24-mediated binding tunes pro-metastatic function of platelets. **A**. Infographics illustrating the generation of shRNA-mediated CD24 knock-down 4T1 cells. **B**. 4T1 shScramble and 4T1 shCD24 cells expression of CD24 predominant isoform (201) mRNA expression calculated using the 2^−ΔΔCt^ method (housekeeping: *GAPDH*). Data are representative of 6 independent experiments. Mann-Whitney test was applied after assessment of gaussian data distribution by Shapiro-Wilk test. **C**. 4T1 shScramble and 4T1 shCD24 cells expression of CD24 minor isoform (202) mRNA expression calculated using the 2^−ΔΔCt^ method (housekeeping: *GAPDH*). Data are representative of 3 independent experiments. Student t-test was applied after assessment of gaussian data distribution by Shapiro-Wilk test. **C**. Flow cytometry assessment of CD24 expression 4T1 shScramble and 4T1 shCD24 cells. Left: Representative histograms of the relative expression levels of CD24-FITC. Isotype signal is represented in black in each graph. Right: Quantification of the mean fluorescent intensity signal for CD24-FITC as fold decrease compared to 4T1 shScramble signal. Data are representative of 10 independent experiments. Mann-Whitney test was applied after assessment of gaussian data distribution by Shapiro-Wilk test. **D**. *In vitro* platelet-TC interaction. Left top: Representative confocal micrographs of platelet-TC binding. In green Wheat Germ Agglutinin (WGA)-iFluor488, in red platelet RAM.1 (Alexa Fluor 647), and in blue nuclei (DAPI). Scale bar = 10µm. Left center: isolated platelet fluorescence channel. Left bottom: binary mask applied on isolated platelet fluorescence channel for circularity analysis. Right top: Quantification of the number of platelets bound per tumor cell. Right bottom: Quantification of platelet circularity. Data are representative of 10 fields of view from 2 independent experiments. Mann-Whitney test (number of platelet/tumor cell) and Student t-test (platelet circularity) were applied after assessment of gaussian data distribution by Shapiro-Wilk test. **E**. Tumor cell clustering upon platelet binding. Left: Infographics illustrating the experimental setting for the TC-platelet cluster formation *in vitro*. Right: Quantifications of the ratio of clusters per single cells and of the cluster area in absence (plasma) or presence (cPRP) of platelet. Ratio of clusters per single cells: Data are representative of 10 fields of view from 2 independent experiments. Student t-test (4T1 shScramble) and Mann-Whitney test (4T1 shCD24) were applied after assessment of gaussian data distribution by Shapiro-Wilk test. Cluster area: Data are representative of between 453 and 498 cells from 2 independent experiments. Mann-Whitney test was applied after assessment of gaussian data distribution by Shapiro-Wilk test. **F**. Tumor cell cluster resistance to anoikis. Left: Infographics illustrating the experimental setting of the anoikis resistance assay. Right: Percentage of Zombie NIR^pos^ (dead) cells as quantified by flow cytometry at 24 hours post clusters formation in absence (plasma) or presence (cPRP) of platelet. Data are representative of 20,000 cells analyzed per conditions from 3 independent experiments. Student t-test was applied after assessment of gaussian data distribution by Shapiro-Wilk test.

### CD24-mediated platelet binding controls lung seeding and colonization and subsequent metastatic outgrowth

Owing to these *in vitro* results, we probed the metastatic potential of 4T1 shScramble and 4T1 shCD24 cells *in vivo*. Building on our previously published results showing that platelet-binding levels differently impact TC lung seeding (8), we first decided to demonstrate that CD24 pro-metastatic function is indeed linked to its platelet-binding function. We depleted platelets with an anti-GPIbα antibody to induce the short thrombocytopenia (TCP) regimen we previously described (8) prior TC injection in syngeneic mouse model of experimental metastasis (**Fig.3A-B**) and longitudinally tracking lung metastatic burden of TC. At the moment of TC seeding and extravasation (up to 3 dpi) (31), platelet depletion impaired extravasation of 4T1 shScramble cells but not 4T1 shCD24 cells (**Fig.3C**). This suggests that CD24-mediated TC-platelet interaction is required for early lung metastatic colonization. When assessing the platelet and CD24 effect on later metastatic outgrowth (14dpi), we further validated that platelets depletion at the time of TC injection massively reduce metastatic outgrowth in both conditions (**Fig.3D**), synergistically blunting metastatic outgrowth in CD24 knock-down condition. Yet, 4T1 shCD24 cells retained a residual metastatic potential in mice with normal platelet counts (**Fig.3D and Fig.S3A-C**), suggesting, as expected, that CD24 is not the sole molecular glue for TC-platelet interaction. To investigate the mechanisms that may underpin the late metastatic outgrowth, metastatic lungs were surgically resected on day 14. Macroscopic inspection of 4T1 shScramble and shCD24 metastatic lung revealed that early TCP significantly reduced the percentage of metastatic area, in agreement with the defect in lung colonization highlighted at 3 dpi and the subsequent lower bioluminescent signal observed at 14dpi (**Fig.3E**). However, these lesions displayed similar surface area and number of Ki67^pos^ cells (**Fig. 3E-F**), suggesting that, while platelet control early lung metastatic colonization through CD24, they do not impact the later proliferation of 4T1 cells during the outgrowth of already-established metastatic foci. Moreover, the lack of difference in Ki67^pos^ cells, consistent with the absence of variations in *in vitro* proliferation (**Fig.S3D**), indicated that our observations were not due to a cell-intrinsic proliferation defect in the 4T1 sublines caused by the disruption of CD24 known role in modulating TC proliferation (32). Altogether, our results suggest that CD24 promotes metastasis by impacting early platelet-dependent seeding and colonization of circulating TC, independently of controlling their proliferation status.

**Figure 3.**
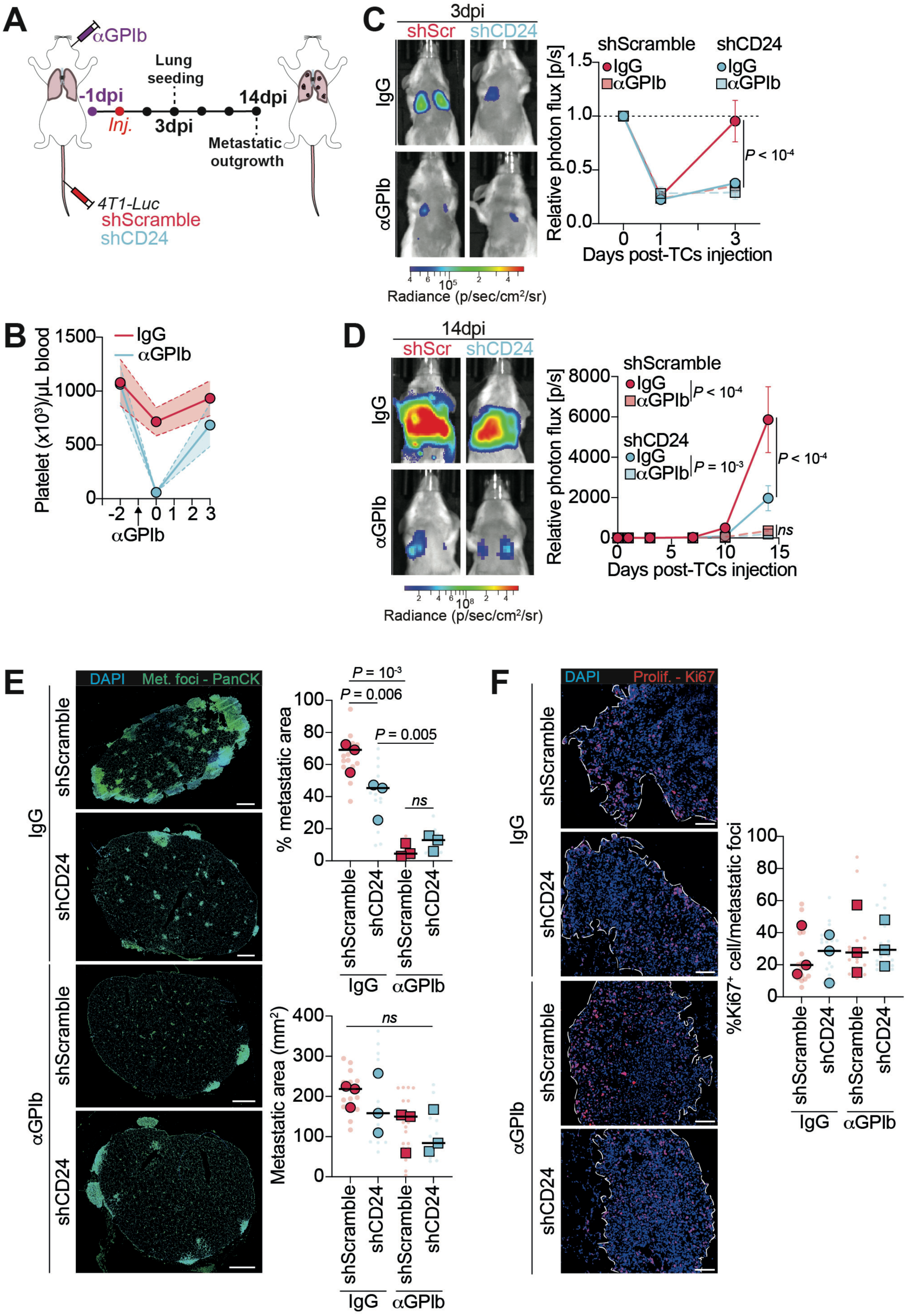
CD24-mediated platelets binding controls early steps of lung colonization and subsequent metastatic outgrowth. **A.** Infographics of short thrombocytopenia regimen (short TCP) within the experimental metastasis model. **B**. Platelet counts in IgG- and αGPIb-treated BALB/c mice. **C-D**. *In vivo* longitudinal Body Luminescence Index (BLI) measurement of 4T1 shScramble or 4T1 shCD24 lung seeding and metastatic outgrowth in normal and TCP mice. **C**. Left: Representative BLI images at 3 dpi (lung seeding). Right: Quantification of the BLI signal represented as the relative photon flux (p/s) on D_0_ signal. Data are representative of 5 mice per group from 1 experiment. Two-way ANOVA corrected with original FDR method of Benjamini-Hochberg was applied. **D**. Left: Representative BLI images at 14 dpi (metastatic outgrowth). Right: Quantification of the BLI signal represented as the relative photon flux (p/s) on D_0_ signal. Data are representative of 5 mice per group from 1 experiment. Two-way ANOVA corrected with original FDR method of Benjamini-Hochberg was applied. **E**. Metastatic burden immunofluorescence staining from OCT-embedded lungs at 14 dpi. Top: Representative images showing (from left to right): DAPI channel (nuclei), PanCK channel (metastatic foci) and merged channels. Scale bar = 1mm. Bottom: Quantification of percentage of metastatic area per slide and metastasis average size (metastatic area). Data are representative of 4 to 6 images per mouse from 3 mice per group and 1 experiment and displayed as mean per mouse (large points) superimposed on the value per single image (small points). One-way ANOVA with original FDR method of Benjamini-Hochberg was applied after assessment of gaussian distribution of the data by Shapiro-Wilk test. **F**. Metastatic tumor cell proliferation immunofluorescence staining from OCT-embedded lungs at 14dpi. Top: Representative images showing, from the left, DAPI channel (nuclei), Ki67 channel (proliferation) and channels merging. Scale bar = 100μm. Bottom: quantification of the percentage of Ki67^pos^ cells per metastatic foci. Data are representative of 4 to 6 images per mouse from 3 mice per group and 1 experiment and displayed as mean per mouse (large points) superimposed on the value per single image (small points). One-way ANOVA with original FDR method of Benjamini-Hochberg was applied after assessment of gaussian distribution of the data by Shapiro-Wilk test.

### Early CD24-mediated tumor-cell-platelet interaction controls later tumor immune microenvironment composition

CD24 knock-down reduced the late metastatic outgrowth of 4T1 cancer cells in the lungs starting at 10 dpi (**Fig.3 and Fig.S3**), suggesting an enhanced adaptive immune response to metastasis. Indeed, we already previously demonstrate the ability of platelet to tune the metastatic TiME and, likely, alter TC fitness and outgrowth (8). Hence, we further explored the mechanisms underlying the reduced metastatic burden in 4T1 shCD24-injected mice using multiparametric flow cytometry. We examined the immune microenvironment of metastasis-bearing lungs of mice with normal platelet counts and found that 4T1 shScramble metastatic nodules induced a more immunosuppressive TiME compared to 4T1 shCD24 nodules (**Fig.4A**). Notably, 4T1 shScramble metastatic outgrowth was associated with a strong enrichment in total neutrophils and in their pro-tumoral Siglec-F^+^ subtype (33) (**Fig.4A**). Conversely, CD24 knock-down rewired lung metastatic microenvironment towards a lymphoid response,as highlighted by the global augmentation in B-, NK- and T cells observed in 4T1 shCD24-tumor bearing mice (**Fig.4A**). Although the proportion of CD4^+^ and CD8^+^ T cells are similar between 4T1 shScramble- and shCD24 lung metastatic lesions, both infiltrated CD4^+^ and CD8^+^ T cells presented an increase in activated populations as per CD44 expression (**Fig.4A**). Remarkably, we also observed a rise in dendritic cells infiltration, further confirming the invigorated T cells activation (**Fig.4A**). Besides, we noticed a significant eosinophils recruitment (**Fig.4A**), which was previously described as promoting the mounting of a lymphocyte-mediated anti-tumor immunity (34). Further analysis of the soluble metastatic lung parenchyma milieu revealed an enrichment in pro-metastatic molecules such as matrix metalloproteases (MMP-3 and MMP-9) (35,36) in the 4T1 shScramble-injected mice whereas 4T1 shCD24 metastatic lesions were associated with an increase in lymphoid and dendritic cells chemoattractants such as CXCL12, CXCL13 and MIP1-α/CCL3 (37–39) (**Fig.S4A**). Importantly, we found that the TiME changes did not correlate with the number of metastases when either 4T1 shScramble and shCD24 lung metastatic lesions were analyzed, suggesting that was unlikely to be the consequence of the impaired colonization and subsequent decreased metastatic burden in the lungs of 4T1 shCD24-injected mice (**Fig.S4B**). Overall, these results suggest that early CD24-mediated TC-platelet interaction controls later metastatic TiME, inducing an immunosuppressive environment. Interestingly, additional analysis of publicly available TNBC databases based on the FUSCC classification revealed that high *CD24* expression level was associated with poor prognosis particularly in the basal-like immune-suppressed (BLIS) TNBC subtypes, further validating its potential role in TiME immunosuppressive phenotype (**Fig.S4C**).

**Figure 4.**
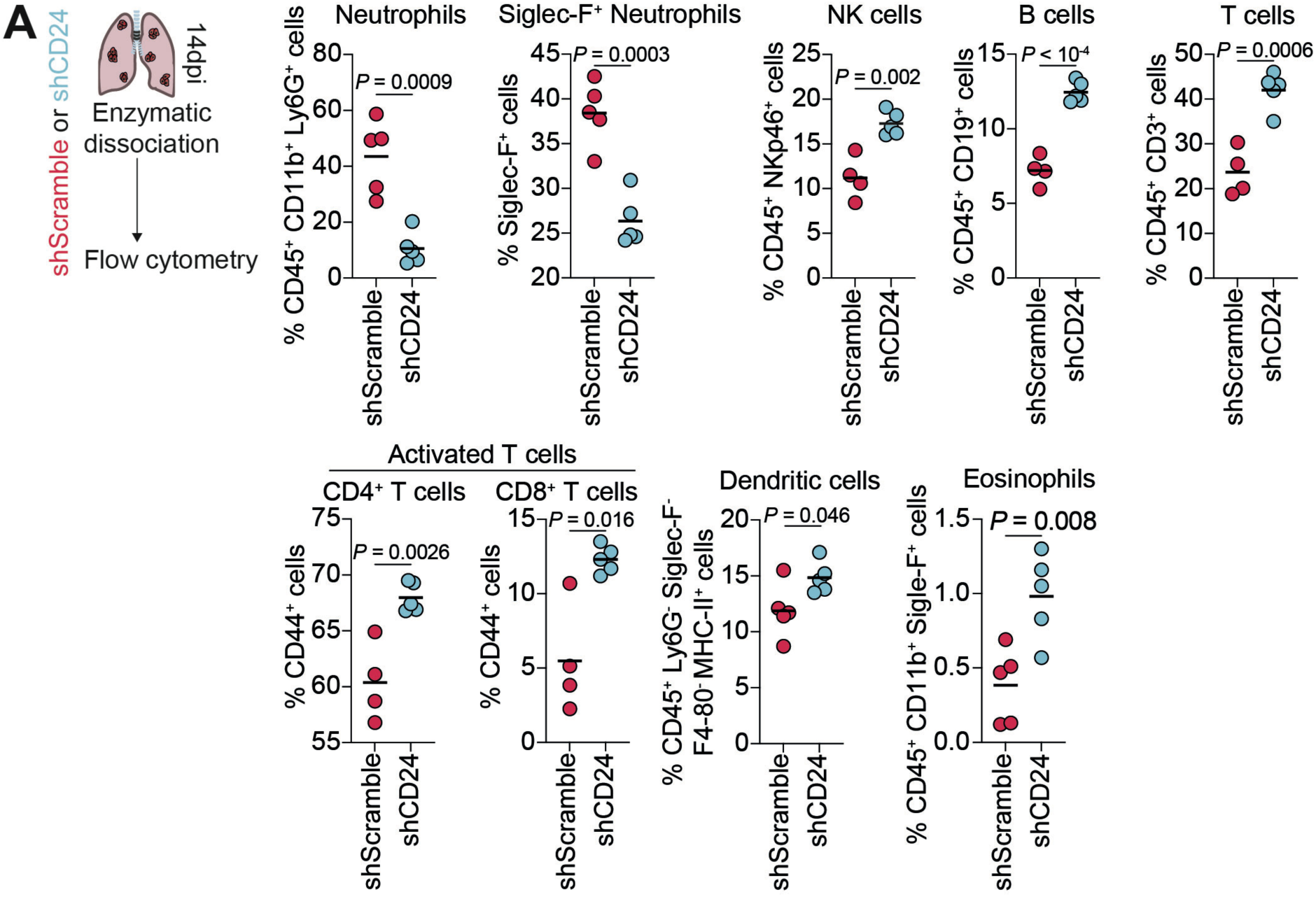
Early CD24-mediated tumor-cell-platelet interaction controls later tumor immune microenvironment composition. **A**. *Ex vivo* multiparametric flow cytometry of immune cells infiltrating mice bearing lung metastases. Left: Infographics describing the experimental scheme. Right: Immune cells population as percentage of their parental cells assessed by flow cytometry analysis at 14dpi. Data are representative of 4 mice per group from 1 experiment. Mann-Whitney (Activated CD8^+^ T cells) or Student t-test (Neutrophils, Siglec-F^+^ neutrophils, NK cells, B cells, T cells, Activated CD4^+^ T cells, Dendritic cells, Eosinophils) were applied after assessment of gaussian data distribution by Shapiro-Wilk test.

## Discussion

This manuscript identifies for the first time CD24-mediated binding of TC to platelet as the determinant of platelets pro-metastatic role. We have previously shown that despite presenting different platelet binding efficacy, TC behave as platelets weak agonist, inducing their aggregation to a similar extend (8). Here, comparing TC presenting different platelet binding abilities, we revealed that they differ in CD24 expression levels. By screening publicly available databases, we revealed that high expression of CD24 in patients correlates with increased malignancy, poorer overall survival as highlighted also in a meta-analysis study (40). We further highlighted that CD24 high expression correlated with a bad prognosis in lymph node-positive TNBC patients, underlining its role in cancer progression. When analyzing our TC collection, we observed that CD24 expression on TC positively correlates with the number of platelets bound to TC and to the metastatic potential of the cell lines, in line with the increased platelet score in lung metastatic vs primary TME. Taking advantage of a shRNA approach, we knocked-down CD24 and we demonstrated that TCs unequal binding to platelets is, in part, mediated by CD24. Of note, CD24-expressing TC displayed bound platelet with a spread morphology whereas platelets binding on CD24-negative TC remained discoid. Upon their activation, platelets express P-selectin at their surface (41) which is essential for their stable binding (20). As CD24 is a ligand of P-selectin, we can hypothesize that while TC similarly activate platelets (8) only the ones expressing CD24 are able to stably bind activated, P-selectin^pos^ platelets. Further invigorating its involvement, we showed that CD24 is required to tune platelets pro-metastatic clustering and resistance to anoikis *in vitro*. As TC-platelet interaction is key to protect TC from shear stress and immune recognition in the bloodstream (4,5), these data suggest that the expression of CD24 might attend this binding hence sustaining the metastatic spreading of CD24^pos^ cells. Prompted by these results, we validated this notion *in vivo* assessing the lung metastatic colonization. In a shortTCP setting, we demonstrated that the early reduction of platelets combined with the depletion of CD24 on TCs completely blunted metastases outgrowth. This additive effect further identifies the early CD24-mediated TC-platelet interaction as a determinant of TC seeding and subsequent metastasis burden. Indeed, 4T1 shCD24 presented a reduced metastatic burden and a significantly diminished number of lung metastatic nodules. Of note, the absence of difference in lung colonization between 4T1 shScramble and 4T1 shCD24 in a shortTCP setting shed a new light on previously published results (14,15), suggesting that platelet but not endothelial P-selectin expression contributes to TC arrest in the lung.

Besides, we found that CD24 expression profoundly impact the metastatic immune microenvironment. Our results are in line with what recently observed in ΔCD24 4T1 primary tumors (42), where authors suggest that the loss of CD24 transform the TME from a “cold” to a “hot” state hence triggering the host response to tumor cells. Furthermore, our data also showed that this immune alteration is linked to a profound rewiring of the cytokine milieu as observed in the cytokine array. A previous study on lung and ovarian cancer already demonstrated that the targeting of CD24, via a blocking antibody, modifies the tumor secretome and reduces tumor growth *in vivo* (43). Whether these effects rely solely on platelets and, moreover, on their early or later recruitment within the tumor bed is still unknown. Indeed, platelets deeply contribute to the reshaping of the TME engaging in a feed-forward communication with TC both through secreted factors or direct exchange (44). In addition, platelets have already been described to infiltrate established primary tumors and activate in contact of TC expressing CD24 (16). Based on these pieces of evidence, we can speculate that CD24-mediated platelets binding influence metastatic outgrowth by *i*) promoting TC colonization and *ii*) rewiring the TiME towards an immunosuppressive one. Altogether, our results suggest that CD24 is, in part, responsible for the differential binding of platelets to TC and the subsequent pro-metastatic function of platelets. In addition, these findings identify CD24 as a potential druggable target to counteract TC-platelet interaction and metastatic progression, avoiding the collateral hemostatic perturbations of classical anti-platelet agents.

## Acknowledgements

We thank all members of JGG and PM teams for their constant discussions on this topic. Image acquisition was performed on the Imaging Platform of the CRBS, PIC-STRA UMS 38, Inserm, Unistra. This work has been directly funded by the French National Institute for Cancer (INCa) grant (PLBIO 2023-173) and by the support of the Ligue Contre le Cancer (labelisation), the Association Ruban Rose, and the SATT Conectus (Strasbourg). VM was supported by a fellowship from the French Ministry of Science (MESRI) and a fourth-year thesis fellowship from the Fondation ARC pour la recherche sur le cancer as well as by the Association Ruban Rose. C.L. and S.M.G.T. are supported by INCa (PLBIO 2023-173). LB is supported by a doctoral fellowship from the Fondation pour la Recherche Médicale.

## Authorship contributions

Conceptualization, V.M., C.L., M.J.G-L., O.L., and J.G.G.; Methodology, O.L.; Investigation, V.M., C.L., C.M., S.M.G.T., L.B., A.F., L.P., and O.L.; Formal Analysis, V.M., C.L., and O.L.; Writing – Original Draft, V.M., C.L., O.L., J.G.G.; Writing – Review & Editing, V.M., C.L., O.L., P.M.; J.G.G.; Supervision, P.M., and J.G.G.; Funding Acquisition, P.M. and J.G.G.

## Material and Methods

### Murine RNA-sequencing analysis

Bulk RNA-sequencing data of 4T1 and 67NR cells obtained by Affymetrix microarray were downloaded from the Gene Expression Omnibus (GEO) database (45) under accession number PRJNA106771. Bulk RNA-sequencing data of 4T1 and B16F10 cells obtained by Illumina HiSeq 2000 were downloaded from the European Nucleotide Archive (ENA) at EMBL-EBI (https://www.ebi.ac.uk/ena/browser/search) under the accession numbers PRJEB5797 (4T1 cells) and PRJEB5299 (B16F10 cells). Sequence reads were mapped to *Mus musculus* mm10 using STAR to obtain a BAM (Binary Alignment Map) file. An abundance matrix was generated based on read counts identified by FeatureCounts.

### Human tumor bulk RNA-sequencing analysis

#### Gene expression levels and survival analysis

RNA-sequencing data regarding expression levels for *CD24* and *PODXL* from human tumors and matched healthy tissues were downloaded as log_2_ (normalized counts +1) values from UCSC Xena (46) (https://xenabrowser.net) with the query “TCGA Breast Cancer (BRCA)”. To analyze CD24 expression during TNBC progression, patients were classified according to their lymph node positivity in the Miller cohort (47) and the TCGA BRCA dataset. For global survival analysis, patients were stratified into high *versus* low *CD24* or *PODXL* expression using their median expression values. For cancer progression survival and progression-free analyses, patients were divided in two groups (LN^pos^ and LN^neg^) based on their lymph node positivity and stratified into high *versus* low *CD24* expression using their median expression values. For survival analyses of TNBC subtypes, data were retrieved using the Pronostic module of bc-GenExMiner (https://bcgenex.ico.unicancer.fr/) (48). Patient data extracted from GSE33926, GSE12276, GSE19615, GSE21653, GSE76124, GSE58812, GSE83937 were categorized into BLIS, BLIA, LAR and MLIA subtypes and stratified into high *versus* low *CD24* expression using their median expression values. Kaplan-Meier plots were generated and analyzed using Prism 9.0.

#### Tumor microenvironment landscape assessment

Bulk RNA-sequencing data of primary breast cancer and matched lung metastatic lesions obtained by Illumina HiSeq 2000 were downloaded from the Gene Expression Omnibus (GEO) database (45) under accession number PRJNA794830. Sequence reads were mapped to *Homo sapiens* hg19 using STAR to obtain a BAM (Binary Alignment Map) file. An abundance matrix was generated based on read counts identified by HTSeq-count. Normalized bulk RNA-sequencing data were used as an input in xCell (http://xCell.ucsf.edu/) (49) to quantify relative cell type abundance within primary and metastatic TME.

#### Genes correlation

The Spearman correlation coefficient ρ was calculated with the Gene_Corr module of TIMER2.0 (http://timer.comp-genomics.org/) (50) to evaluate the co-expression patterns between *GP6* and the following genes: *CD24*, *CD44*, *SELPLG*, *PODXL*, *PDPN*, *ITGAV*, *ITGB3*, *CD97*, and *LGALS3* in the TNBC subset of the TCGA BRCA cohort.

### Single-cell RNA-sequencing analysis

Single-cell RNA-sequencing data from (24) were accessed on the Broad Institute Single Cell portal (https://singlecell.broadinstitute.org/single_cell) with the query “*CD24*” and “*PODXL*”. Mean gene expressions at the single cell level for each annotated cell types and percentage of expressing cells were extracted from the dot plot representation.

### Cell culture

The mouse breast cancer cell lines 4T1, 67NR, AT3, D2A1 and the mouse melanoma cell line B16F10, were cultured under standard conditions (37°C, 5% CO_2_) using RPMI-1640, supplemented with 10% FBS and 1% penicillin-streptomycin solution. The human embryonic kidney HEK293T cell line was cultured under standard conditions (37°C, 5% CO_2_) using DMEM, supplemented with 10% FBS and 1% penicillin-streptomycin solution. Cell viability *in vitro* was assayed before experiment by a Countess 3^®^ automated cell counter (ThermoFisher).

### Generation of 4T1 sub-lines

#### Plasmid construction

Control or CD24 double stranded oligos targeting sequences (**Table S1**) were inserted into the AgeI to EcoRI sites of pLKO.1 vector (Addgene #10878).

#### Lentivirus production

HEK293T cells were seeded in a 6 well plate and transfected using JetPrime reagent (Polyplus, #101000027) with either pLKO.1-shScramble or pLKO.1-shCD24, together with pLP1, pLP2 and pLP3/VSV-G lentiviral packaging plasmids (Invitrogen). Two days later, the viral lentivirus-containing supernatant was filtered through a 0.22µm filter and the viral particles were precipitated using the Lenti-X concentrator (Clontech, #PT4421-2). After centrifugation at 1500*g* during 45 min at 4°C, the high-titer virus-containing pellets were resuspended in 1 mL of PBS and aliquoted before storage at −80°C.

#### 4T1 cells transduction

4T1-Luc cells were seeded in a 6 well plate and transduced with one aliquot (100µL) of pLKO.1-shScramble or pLKO.1-shCD24 lentivirus, in the presence of 5µg/mL of polybrene (Sigma Aldrich, #H9268). Two days later, transduced 4T1-Luc cells were selected with blasticidin at 5 µg/mL (Sigma Aldrich, #15205-100MG), to generate the stable 4T1-Luc-shScramble or shCD24 sublines.

### RT-qPCR analysis

To measure CD24 predominant (201) and minor (202) isoforms expression, total RNA was extracted using TRI Reagent (Molecular Research Center). Next, complementary DNA (cDNA) was prepared with the High-Capacity cDNA Reverse Transcription Kit (Applied Biosystems #4368814), according to the manufacturer’s instructions. Semi-quantitative PCR analysis on each biological sample was performed using technical replicates with the FastStart Universal SYBR Green Probe Master (ROX) mix for CD24 isoforms or the TaqMan system for the GAPDH housekeeping gene on a 7500 Real-Time PCR system. The cDNA concentration of target genes was normalized by amplification of GAPDH and fold changes in gene expression were obtained using the 2^-ΔΔCt^ method. Custom primers for CD24 isoforms were CD24_201 (Forward: TCTCTTTGGAATTGAACGTTTTG; Reverse: TGCCTGGCACATGTTAATTACT) and CD24_202 (Forward: TCGTCTTCCCCTTCTCAGG; Reverse: TCTGGTGGTAGCGTTACTTGG). TaqMan probe for GAPDH (Mm99999915_g1).

### CD24 surface expression

Murine tumor cells (4T1 native, 4T1 shScramble, 4T1 shCD24, 67NR or B16F10) were washed with EDTA 0.48 mM (Versene, Gibco) and gently detached using a 0.25% trypsin-0.02% EDTA solution (Gibco). After a wash in media containing 10% FBS, cells were resuspended in PBS containing 2% FBS and 2mM EDTA. Murine tumor cells were stained with FITC anti-CD24 (1:200, BioLegend, clone M1/69) or its control isotype (Rat IgG2bk, eBioscience, clone eB149/10H5) for 15 minutes at 4°C. Samples were acquired with Attune NxT (Invitrogen) flow cytometer and data were analyzed using FlowJo™ v10 Software (ThreeStar).

### Citrated platelet-rich plasma preparation

Murine blood was collected by aortic puncture in sodium citrate (0.315%). Citrated platelet-rich plasma (cPRP) was obtained by centrifuging whole blood in microtubes during 1 min at 1,900*g* at 37°C. The platelet count was adjusted to 300×10^3^/µl with citrated platelet-free plasma (cPFP). Mice were anesthetized (intra-peritoneal, *i.p.*, ketamine 100mg/kg, xylazine 10mg/kg) during the procedure and sacrificed by cervical dislocation at the end.

### Platelet binding to tumor cell

For TC-platelet interaction, 6,000 tumor cells (4T1 native, 4T1 shScramble, 4T1 shCD24, 67NR or B16F10) were seeded per well on a 12 wells chamber and let to adhere overnight in complete medium. The following day, cPRP was added to each well at the ratio of 500 platelet/cell and let to interact for 30 minutes at 37°C 5% CO_2_ in PBS. During the last two minutes of interaction, Wheat Germ Agglutinin-iFluor 488 (10µg/mL) was added. After three PBS washes, cells were fixed in PFA 0.5% in PBS for 20 minutes at room temperature. Cells were washed three times and stained with Alexa Fluor 647 coupled anti-RAM.1 antibodies (2μg/mL) for 1 hour at room temperature. Slides were then washed and mounted in Fluoromount-G™ with DAPI. Images were acquired with 60x/water objective at Olympus Spinning disk confocal microscope. A z-stack covering the full height of the tumor cell was acquired to ensure that only platelets in direct contact with the cells were imaged. Maximum intensity z-projections were generated using Fiji 2.15.1, and the number platelet engaging in interactions with tumor cell was manually quantified. Final data processing was performed in Excel 6.0 and GraphPad Prism 9.5 for statistical analysis.

### Platelet circularity

To assess platelet circularity, images acquired for TC-platelet interaction were processed with Fiji 2.15.1. Fluorescence channels were split and platelet fluorescence, corresponding to anti-RAM.1 antibody signal, was segmented using the ‘Default’ thresholding method. Then, segmented platelet circularity was measure using the ‘Analyzed particles’ plugin. Data were post-processed in Excel 6.0 and GraphPad Prism 9.5 for statistical analysis.

### Tumor cell clusters formation upon platelet binding

To form TC clusters in vitro, 2.10^6^ 4T1 shScramble or shCD24 cells in suspension were co-incubated with 2.10^7^ murine cPRP for 2 hours at 37°C 5%CO_2_ on a tilting tray. Cells were then washed with serum-free medium, and clusters were imaged on an inverted microscope. Macroscopic RGB images with identical zooming were loaded on ImageJ and clusters’ area was measured after polygon selection definition.

### Anoikis resistance assay upon platelet binding

To determine the response to anoikis clustered cells were plated at 500,000 cells/mL in 6-well ultra-low adherence plate in complete culture medium for 24h. At the end of the incubation, cells were harvested and stained with Zombie NIR (1:1,000; Biolegend, #423105) for 15 minutes at RT in the dark, prior to wash and signals’ acquisition on an Attune NxT (Invitrogen) flow cytometer (20,000 events/conditions). Data were analyzed using FlowJo™ v10 Software (TreeStar).

### Mice

All animals were housed and handled at INSERM (mouse facility agreement number: #C67-482-33) according to the guidelines of INSERM and the ethical committee of Alsace (CREMEAS), following French and European Union animal welfare guidelines (Directive 2010/63/EU on the protection of animals used for scientific purposes). All procedures were performed in accordance with French and European Union animal welfare guidelines and supervised by local ethics committee under the following APAFIS authorization #37433-2022052016445806.

Eight-week-old female wild-type immunocompetent BALB/c mice (028BALB/C; Charles River) were used in all experiments. Animals were housed in pathogen-free conditions with food and water *ad libitum* and appropriate enrichment (sterile pulp paper and coarsely litter). Mice were monitored daily, and terminally sick animals were euthanized under an approved protocol.

### Experimental pulmonary metastasis assay and bioluminescent imaging

For the experimental pulmonary metastasis experiment, subconfluent 4T1-Luc (shScramble or shCD24) were washed with EDTA 0.48 mM (Versene, Gibco), gently detached using a 0.25% trypsin-0.02% EDTA solution (Gibco), washed in media containing 10% FBS, resuspended at 1.5 x 10^6^ TC/mL in serum-free media, filtered through a 40 µm mesh (Falcon) and kept on ice until injection. Viability (>85% prior injection) was determined by Trypan Blue exclusion. Tumor cells (100µL, containing 150,000 cells) were injected through the lateral tail vein (intra-venously, *i.v*.) with a 25-gauge needle. *In vivo* imaging was performed shortly after to establish initial lung seeding (15 minutes post-injection) by *i.p.* injection of D-luciferin solution (150mg/kg) and subsequent signal acquisition in isofluorane (Isoflo, Zeotis)-anesthetized mice with an IVIS Lumina III (Perkin Elmer) imaging system. The rate of total light emission of the lung metastatic area was analyzed using the Living Image software (Perkin Elmer) and expressed as numbers of photons emitted per second (p/s). Metastatic outgrowth was longitudinally followed by subsequent imaging at 1-, 3-, 7-, 10- and 14-days post-injection (dpi). For these later timepoints, the relative total light emission of the lung metastatic area compared to day 0 was calculated and represented.

### Platelets depletion

Severe thrombocytopenia was induced by retro-orbital *i.v.* injection of 2 mg/kg of in-house-produced αGPIb antibody (RAM.6) (51) or IgG isotype control per mouse. RAM.6 or IgG were administrated the day before TC injection to induce a short thrombocytopenia (shortTCP). To validate platelet depletion, murine whole blood was collected into EDTA (6 mM) after severing the mouse tail. Platelet count and size were determined in an automatic cell counter (Element HT5, Heska).

### Lung immune microenvironment analysis by flow cytometry

Single cell suspension was obtained from freshly harvested left lung lobes. Tissue was cut into approximately 2mm^3^ pieces using a scalpel, then enzymatically digested with the mouse tumor dissociation kit (MiltenyiBiotec, #130-096-730) at 37°C for 42 minutes using the gentleMACS Octo Dissociator (MiltenyiBiotec) according to manufacturer’s protocol. Cell suspension was filtered through a 50µm mesh and red blood cells were lysed in ACK Buffer [150mM NH_4_Cl, 10mM KHCO_3_, 0.1mM Na_2_EDTA]. Non-viable cells were stained with Fixable Viability Dye-eFluor™780 (1:1,000; eBioscience, #65-0865-14) for 15 minutes at room temperature in the dark. Aspecific antibodies’ binding was minimized with anti-CD32/CD16 blocking antibody (TruStain FcX 1:50, Biolegend #101320) for 20 minutes at 4°C. Primary conjugated (**Table S1**) were incubated for 15 minutes at 4°C. Samples were acquired with Attune NxT (Invitrogen) flow cytometer and data were analyzed using FlowJo™ v10 Software (ThreeStar). Gating strategies are displayed in Fig.S5.

### Tissue array

Tissue homogenates were prepared according to manufacturer’s instructions (Mouse angiogenesis kit (#ARY0015) and Cytokine panel kit (#ARY006)). Briefly, lungs were collected and snap frozen in liquid nitrogen at time of sacrifice. Frozen tissues were dissociated with Bead Ruptor 4 dissociator (OMNI International) using 1.5mm Triple-Pure High Impact Zirconium beads (Benchmark) in PBS containing Protease/Phosphatase inhibitors (Pierce). Triton X-100 (Sigma) at the 1% final concentration was added at the end of the process. Supernatants were obtained after spinning the homogenates at 10,000*g* for 5 minutes at frozen at −80°C until analysis. Three-hundred µg of proteins (quantified by Bradford assay – BioRad) were applied per membrane following manufacturer’s instructions and incubated overnight at 4°C in end-to-end shaking. Membranes were revealed using iBright1500 (Invitrogen) after sequential exposure. Images were analyzed in Fiji 6.0. and data were then post-processed in Excel 6.0 and GraphPad Prism 9.5 for statistical analysis.

### Tissue histology and immunofluorescence

At the sacrifice, lungs were inflated in OCT (Tissue freezing media, Leica) before embedding and freezing at −80°C. Slices (7μm) were cut on SuperFrost UltraPlus slides (Thermo Fisher Scientific). After rehydration in PBS, slices were permeabilized in with 0.3% Triton X-100 in PBS for two consecutive 10 minutes incubations. Aspecific antibody binding was minimized by incubating for 1 hour in a Histoblock solution (3% bovine serum albumin-BSA, 5% non-heat inactivated FBS, 20mM MgCl_2_ and 0.3% Triton X-100, in PBS). Immunostaining was performed by diluting the primary antibodies (**Table S1**) in the Histoblock solution and incubating them overnight at 4°C. After two washes in PBS, the appropriate secondary antibodies (**Table S1**) were incubated for 1 hour at room temperature in BSA 5% before mounting (Fluoromount/DAPI). Background and nonspecific staining controls were used. Slides were imaged with a 20X/NA0.8 air-objective on a Slide Scanner microscope VS200 (Olympus). Images were processed using QuPath 0.5.0 (52) and analyzed using consecutive thresholding methods and/or positive cell detection plug-in (Ki67^+^ analysis). For the analyses of metastatic areas, metastases were manually annotated as area presenting high PanCK staining. Data were then post-processed in Excel 6.0 and GraphPad Prism 9.5 for statistical analysis.

### Statistics

Statistical analysis was performed with GraphPad Prism 9.5. The normal distribution of the data sets was assessed by the Shapiro–Wilk normality test. According to the number of data sets compared, Student’s t-test test or One-way ANOVA test with original FDR method of Benjamini-Hochberg were applied (*p<0.05; **p<0.01, ***p<0.001, ****p<0.0001). In all cases, the α-level was set at 0.05. All the data in graphs were presented as median ± standard deviation.

### Data Availability

All data supporting the findings of this study are available from the corresponding authors on reasonable request.

---

**Supplementary Figure 1 related to Figure 1.**
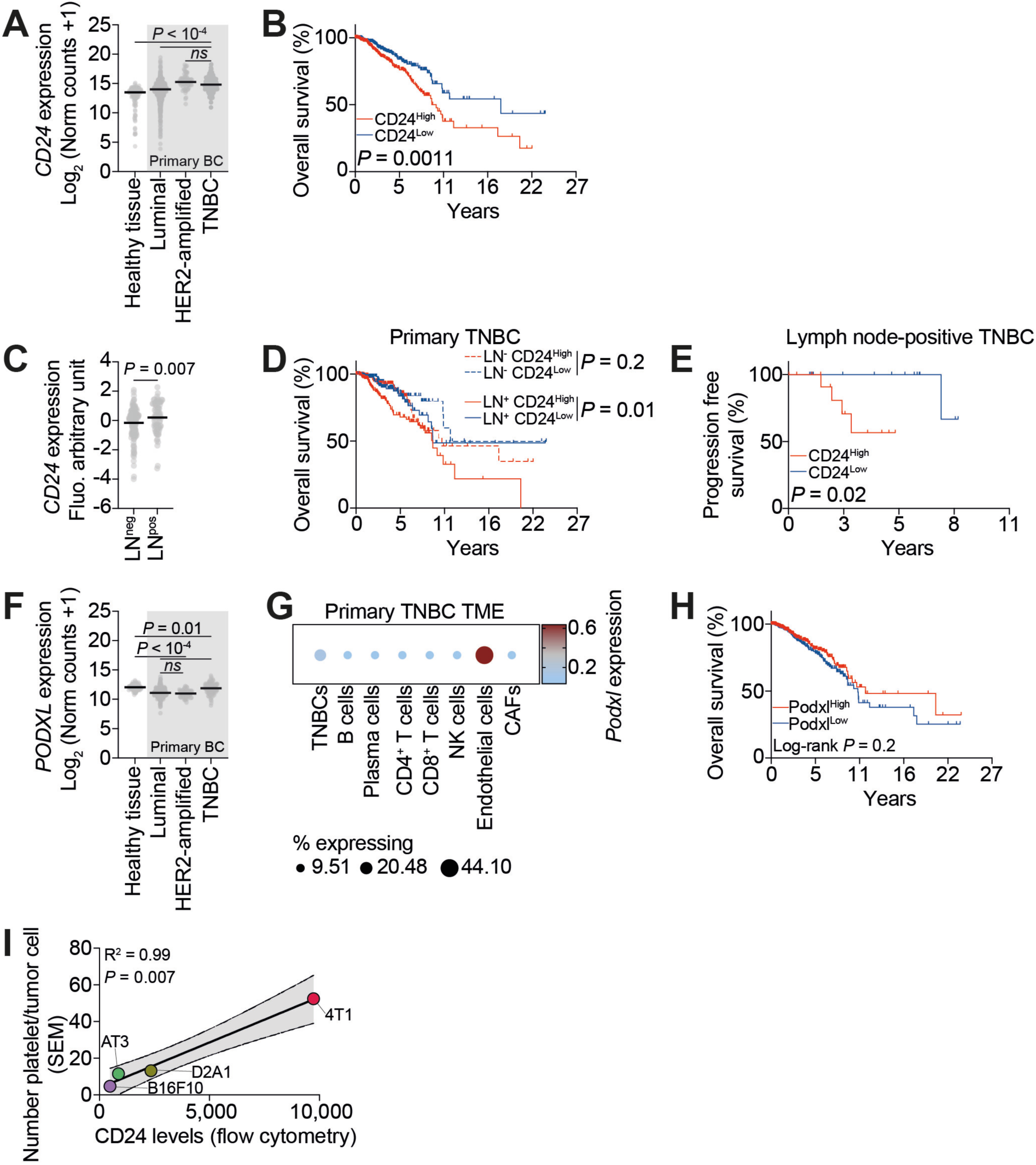
**A**. *CD24* gene expression in healthy and neoplastic breast tissues, with primary breast cancer classified according to the main molecular subtypes. Kruskal-Wallis test with original FDR method of Benjamini-Hochberg was applied after assessment of gaussian distribution of the data by Shapiro-Wilk test. **B**. Overall survival percentage for breast cancer patients (*n* = 1080) with *CD24*^high^ versus *CD24*^low^ expression as defined by median. Two-sided *P* value computed by a log-rank (Mantel-Cox) test. **C**. *CD24* gene expression (Affymetrix array) comparing patients without (LN^neg^, *n* = 158) or with (LN^pos^, *n* = 84) lymph node invasion. Mann-Whitney test was applied after assessment of gaussian data distribution by Shapiro-Wilk test. **D**. Overall survival percentage for LN^pos^ (*n* = 405) and LN^neg^ (*n* = 384) breast cancer patients with *CD24*^high^ versus *CD24*^low^ expression as defined by median. Two-sided *P* value computed by a log-rank (Mantel-Cox) test. **E**. Progression-free survival percentage for LN^pos^ (*n* = 30) breast cancer patients with *CD24*^high^ versus *CD24*^low^ expression as defined by median. Two-sided *P* value computed by a log-rank (Mantel-Cox) test. **F**. Podocalyxin (*PODXL*) gene expression in healthy and neoplastic breast tissues, with primary breast cancer classified according to the main molecular subtypes. Kruskal-Wallis test with original FDR method of Benjamini-Hochberg was applied after assessment of gaussian distribution of the data by Shapiro-Wilk test. **G**. Bubble plot displaying *PODXL* gene expression at the single-cell level across different cell types in the triple-negative breast cancer (TNBC) primary tumor microenvironment (TME). Bubble size represents percentage of *PODX*L-expressing cells and color represent mean *PODXL* expression. CAFs: Cancer Associated Fibroblasts. **H**. Overall survival percentage for breast cancer patients (*n* = 1080) with *PODXL*^high^ versus *PODXL*^low^ expression as defined by median. Two-sided *P* value computed by a log-rank (Mantel-Cox) test. **I**. Pearson correlation between the number of platelets bound to tumor cell and mean CD24 expression levels on tumor cells – 4 xy pairs; R^2^ = 0.99 – 95% confidence interval [0.71-0.99]. Number of platelets interacting with tumor cell were quantified by scanning electron microscopy in (8). Mean CD24 expression levels were quantified by flow cytometry from Fig.1G.

**Supplementary Figure 2 related to Figure 2.**
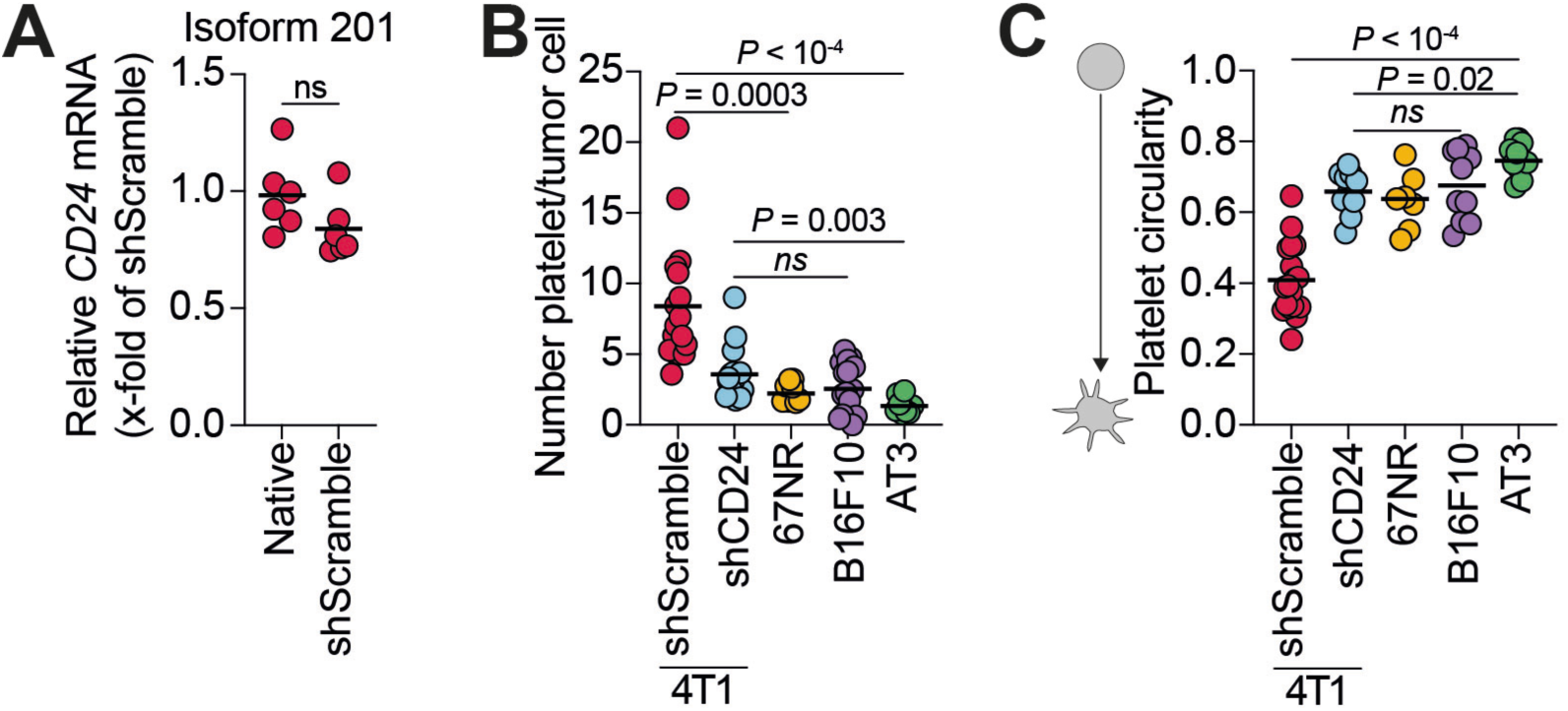
**A**. 4T1 native and 4T1 shScramble cells expression of CD24 predominant isoform (201) mRNA expression calculated using the 2^−ΔΔCt^ method (housekeeping: *GAPDH*). Data are representative of 6 independent experiments. Mann-Whitney test was applied after assessment of gaussian data distribution by Shapiro-Wilk test. **B**. Number of platelets bound per tumor cell as quantified by immunofluorescence staining. Data are representative of 10 fields of view from 2 independent experiments. Kruskal-Wallis test with original FDR method of Benjamini-Hochberg was applied after assessment of gaussian distribution by Shapiro-Wilk test. **C**. Platelet circularity as measured by immunofluorescence staining. Data are representative of 10 fields of view from 2 independent experiments. One-way ANOVA with original FDR method of Benjamini-Hochberg was applied after assessment of gaussian distribution by Shapiro-Wilk test.

**Supplementary Figure 3 related to Figure 3.**
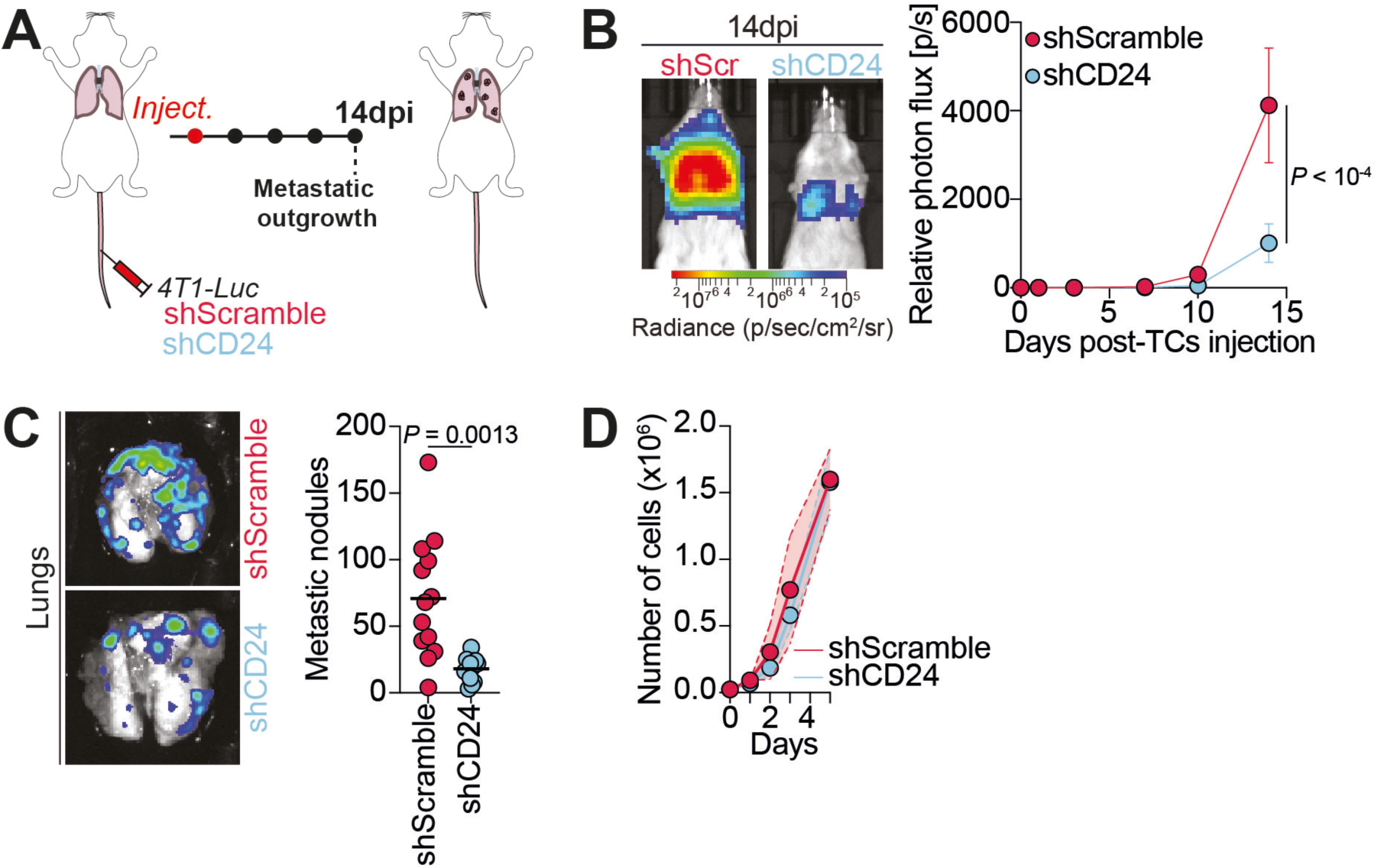
**A.** Infographics of the experimental metastasis model. **B**. *In vivo* longitudinal Body Luminescence Index (BLI) measurement. Left: Representative BLI images at 14 dpi (metastatic outgrowth). Right: Quantification of the BLI signal represented as the relative photon flux (p/s) on D_0_ signal. Data are representative of 10 mice from 2 independent experiments. Two-way ANOVA corrected with original FDR method of Benjamini-Hochberg was applied. **C**. Lungs macroscopic nodules count. Left: Representative BLI images of *ex vivo* lungs at 14 dpi. Right: Quantification of the number of metastatic nodules per lung. Data are representative of 13 mice from 3 independent experiments. Welch’s t-test was applied after assessment of gaussian data distribution by Shapiro-Wilk test. **D**. Number of 4T1 shScramble and 4T1 shCD24 cells per culture days as measured by automatic counter. Data are presentative from 3 independent experiments. Two-way ANOVA corrected with original FDR method of Benjamini-Hochberg was applied.

**Supplementary Figure 4 related to Figure 4.**
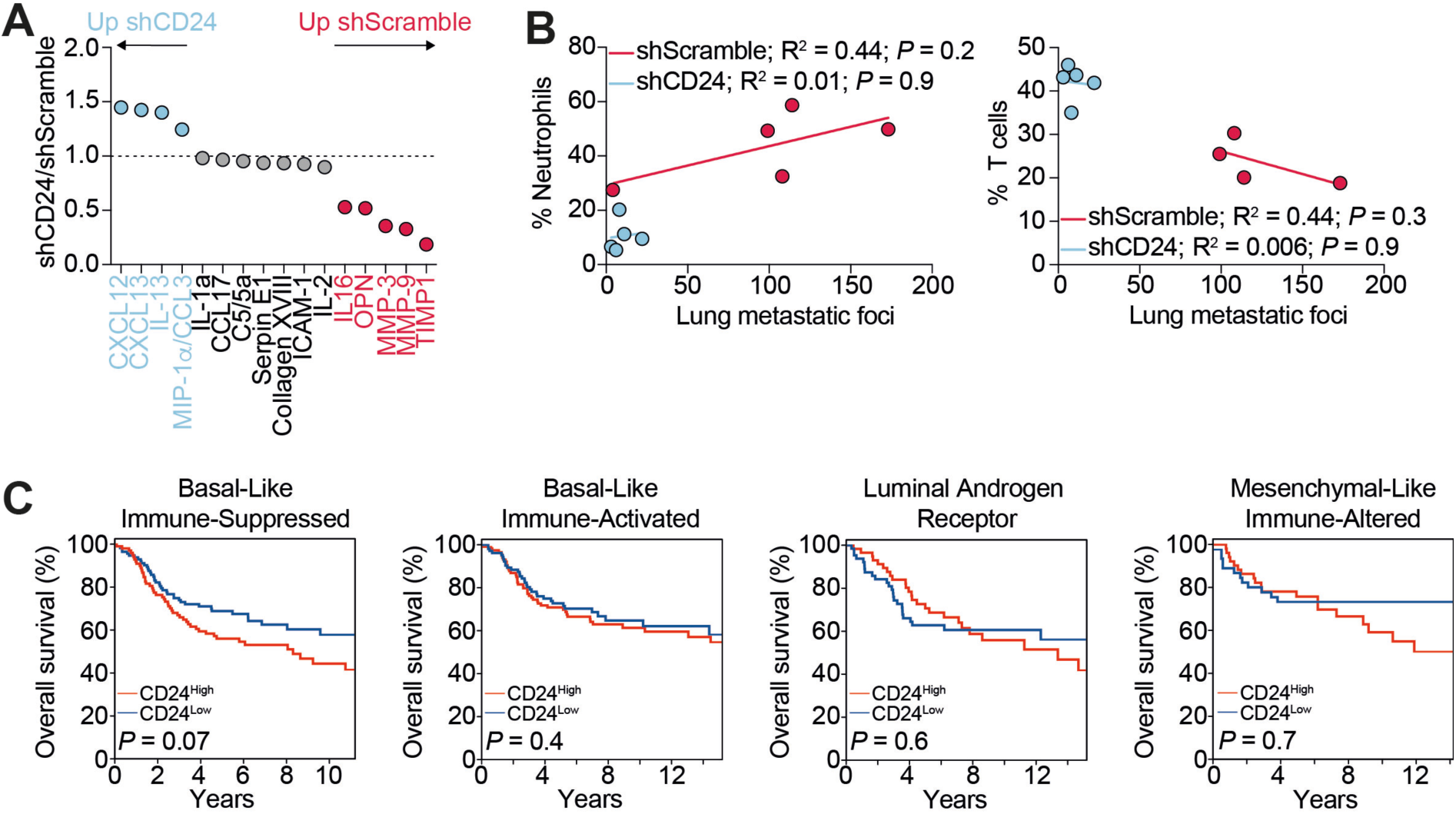
**A**. Tissue array analysis. Relative mean signal per cytokine/chemokine is represented. Data are representative of 2 mice per group from 1 experiment. **B**. Relationship between metastatic burden and infiltration of neutrophils and T cells in the lungs of 4T1 shScramble- (*n* = 5) or 4T1 shCD24- (*n* = 5) bearing mice at 14 dpi. Pearson correlation was calculated on 5 xy pairs. **C**. Overall survival percentage for TNBC breast cancer patients (*n* = 1,015) with different subtypes based on *CD24*^high^ versus *CD24*^low^ expression as defined by median. Two-sided *P* value computed by a log-rank (Mantel-Cox) test.

